# HiChIP: Efficient and sensitive analysis of protein-directed genome architecture

**DOI:** 10.1101/073619

**Authors:** Maxwell R. Mumbach, Adam J. Rubin, Ryan A. Flynn, Chao Dai, Paul A. Khavari, William J. Greenleaf, Howard Y. Chang

## Abstract

Genome conformation is central to gene control but challenging to interrogate. Here we present HiChIP, a protein-centric chromatin conformation method. HiChIP improves the yield of conformation-informative reads by over 10-fold and lowers input requirement over 100-fold relative to ChIA-PET. HiChIP of cohesin reveals multi-scale genome architecture with greater signal to background than *in situ* Hi-C. Thus, HiChIP adds to the toolbox of 3D genome structure and regulation for diverse biomedical applications.

## Main Text

Protein factors help to guide gene regulatory circuits inside living cells. Many techniques exist to map the 1-D landscape of protein binding to the genome, however there are fewer methods for understanding 3-D features. Chromosome conformation capture (3C) coupled to sequencing (Hi-C) has been transformative in our ability to understand, at high resolution, the architecture of the genome^1–5^. However, because Hi-C samples all possible proximity ligations in the genome, very deep sequencing is required to fully identify chromatin architectural features. To achieve enhanced specificity, enrichment strategies have been developed to target factor-directed or locus-directed chromatin interactions: Chromatin Interaction Analysis by Paired-End Tag Sequencing (ChIA-PET) and Capture-C^6–8^, respectively. ChIA-PET combines chromatin immunoprecipitation (ChIP) with 3C, producing a directed view of long-range contacts associated with a protein factor of interest^9^. Despite recent advances, ChIA-PET still requires hundreds of millions of cells per experiment and results in a small fraction of informative reads for a given sequencing depth^10^ **(Supplementary Table 1).** Therefore, improved methods for mapping factor-directed chromatin conformation are needed.

To address these problems, we developed HiChIP, a method that leverages principles of *in situ* Hi-C^5^ and transposase-mediated on-bead library construction^11^ (Fig. 1a). In HiChIP, long-range DNA contacts are first established *in situ* in the nucleus prior to lysis, minimizing possible false-positive interactions^12^ and greatly improving DNA contact capture efficiency. ChIP is then performed on the contact library, directly capturing long-range interactions associated with a protein of interest. Paired-end sequencing then identifies two distantly located segments of the genome from one fragment, indicating that the factor of interest was associated with the long-range interaction. The HiChIP protocol is robust, reproducible, and can be completed in as few as two days (Fig. 1a).

**Figure 1.**
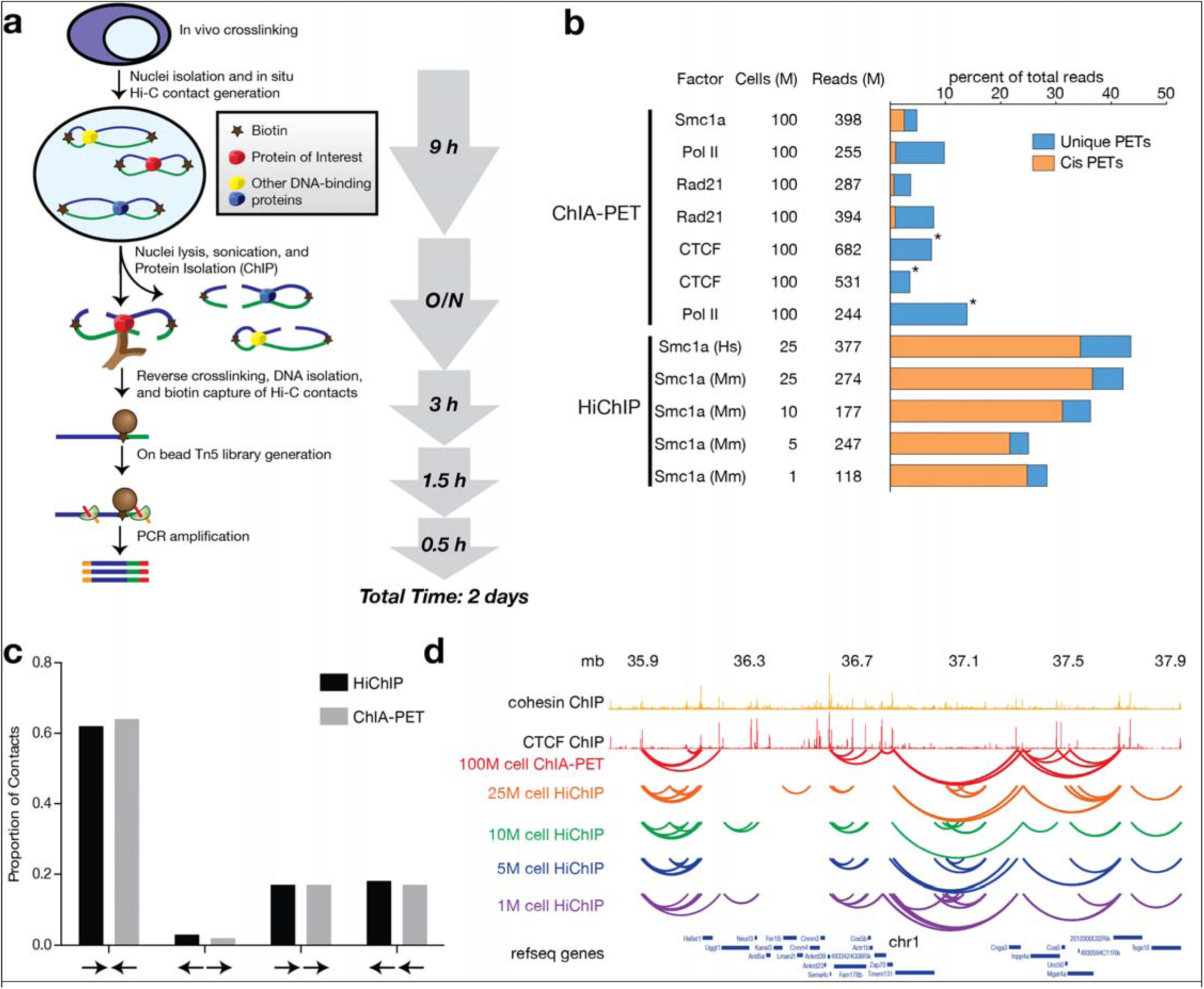
HiChIP is a rapid and sensitive method to discover protein-centric *in situ* chromatin loops. **a,** Schematic of HiChIP method. Briefly, cells are crosslinked and *in situ* contacts are generated. Nuclei are sonicated and ChIP is performed, enriching for contacts associated with a protein of interest. Biotinylated contacts are then captured, and sequencing libraries are generated using Tn5 transposase on the capture beads. The entire HiChIP protocol can be completed in as few as two days. **b,** Efficiency of information obtained from sequencing for two replicates of HiChIP relative to published ChIA-PET studies. HiChIP maintains a robust increase in efficiency even at cell numbers 100-fold lower than ChIA-PET. * denotes ChIA-PET studies which do not report cis and trans PET distinctions, see **Supplementary Table 1. c,** CTCF motif orientation analysis at Smc1a HiChIP contact anchors in GM12878 cells compared to CTCF Advanced ChIA-PET^7^. d, Smc1a ChIA-PET^13^ and HiChIP peak-filtered contacts in V6.5 mouse embryonic stem cells with differing starting cell number.

We performed HiChIP on the cohesin subunit Smc1a in the human B lymphocyte GM12878 (GM) and v6.5 mouse embryonic stem (mES) cell lines to validate our method in comparison to published ChIA-PET datasets^10,13–15^. We processed our HiChIP data using the HiC-Pro pipeline^16^ to filter our total sequenced reads into informative unique paired-end tags (PETs) **(Supplementary Table 2).** Contact calling is highly sensitive to the number of unique PETs observed at a given sequencing depth, making the unique PET efficiency from total reads an important indicator of method efficiency. Greater than 40% of the total sequenced reads were informative PETs in our HiChIP of Smc1a in the GM and mES cell lines. By comparison, ChIA-PET studies report dramatically lower efficiencies (3-12%) with a greater fraction of reads mapping as trans-interactions, a feature generally associated with false positives^12^ (Fig. 1b, **Supplementary Table 1).** To directly compare HiChIP and ChIA-PET at the level of contact calling confidence, we ran Mango^17^ on our GM Smc1a HiChIP data **(Methods).** We found that approximately 50% of HiChIP contacts were supported by at least 40 PETs, compared to 7% for ChIA-PET **(Supplementary Figure 1a).** Consequently, HiChIP both provides a higher percentage of informative reads from total sequencing depth as well as uses those informative reads to call contacts with greater confidence relative to ChIA-PET.

To first address enrichment specificity, we sought to identify 1D ChIP peaks from our HiChIP data as is done with ChIA-PET. In our GM Smc1a HiChIP dataset, we called a total of 27,697 ChIP peaks using the MACS2^18^ peak caller. 79% of our peaks overlapped with ENCODE Smc3 ChIP peaks; our peak set also represented a stringent high confidence subset of the ENCODE peaks **(Supplementary Figure 1b,c).** Therefore, HiChIP is able to provide confident 1D factor binding information.

We applied the Fit-Hi-C method^19^ to identify contacts in our HiChIP experiments. Contact anchors have been reported to be enriched for CTCF and cohesin binding. Furthermore, CTCF motifs in anchors are commonly found in a convergent orientation^5^. We assessed the CTCF motif orientation at contact anchors from our GM Smc1a HiChIP contacts as well as published Advanced ChIA-PET contacts (Fig. 1C). We reproduce the predicted and previously observed distribution of motif orientations, strongly enriching for convergent CTCF motifs.

One of the major limitations of ChIA-PET is the large amount of starting material required, generally at least 100 million cells^9,10,13–15^. To identify minimum input levels for HiChIP, we generated Smc1a HiChIP maps starting from 10, 5, and 1 million mES cells. We found that HiChIP at these lower cell inputs provided comparable fractions and absolute numbers of informative reads to ChIA-PET studies with 100-fold more starting material, despite sequencing our HiChIP libraries to a much lower depth (Fig. 1b, **Supplementary Table 1).** While data from lower cell number samples had fewer PETs (Fig. 1b), we found that contact calls retained a high degree of overlap with our 25M cell samples, underlining the robustness of the data from limited starting material (Fig. 1d).

The *in situ* Hi-C libraries sequenced to approximately 6.5 billion reads in GM cells represents a gold standard of global chromatin conformation^5^. Given the sensitivity and efficiency gains of HiChIP compared to ChIA-PET, we next compared HiChIP to *in situ* Hi-C. We first examined interaction maps of Smc1a HiChIP at an example locus on chromosome 8 from the *in situ* Hi-C study and found that HiChIP identifies chromatin features originally found in Hi-C across several length scales (Fig. 2a).

We next aimed to compare our loop set globally with loops derived from *in situ* Hi-C^5^. We processed our GM Smc1a HiChIP dataset with Juicer^5,20,21^ using the same parameters as the *in situ* Hi-C dataset **(Supplementary Table 3).** We obtained a highly overlapping set of loops as compared to *in situ* Hi-C (Fig. 2b). This degree of overlap was similar to that of the biological replicate data between the primary and replicate *in situ* Hi-C experiments. **(Supplementary Fig. 2a).** Furthermore, direct comparison of the reads supporting the union set of loops *(in situ* Hi-C and HiChIP) revealed a strong correlation between the two experimental strategies (Fig. 2c). Importantly, we identified a similar number of loops (10,255 compared to 9,448) with 10-fold less sequencing.

**Figure 2.**
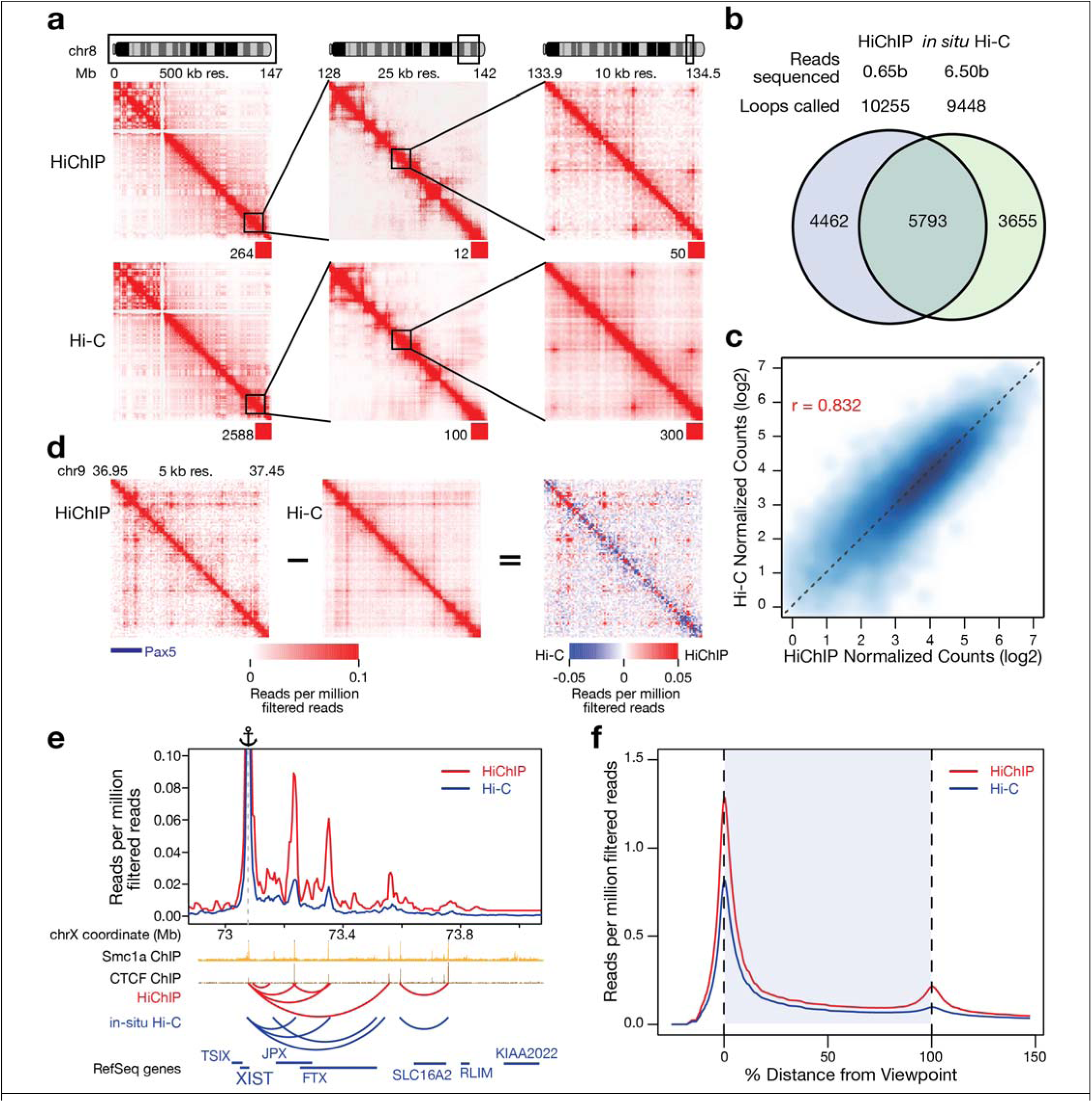
HiChIP of cohesin provides a high signal to background chromatin interaction map. **a,** GM12878 Smc1a HiChIP raw interaction maps of an *in situ* Hi-C example locus on chromosome 8. Numbers below the interaction maps correspond to maximal signal in the matrix. **b,** Overlap of global loop calls from GM12878 Smc1a HiChIP and *in situ* Hi-C primary + replicate library sets. **c,** Reproducibility of loop calls in GM12878 Smc1a HiChIP and *situ* Hi-C primary + replicate library sets. d, Read normalized *in situ* Hi-C and HiChIP raw maps at the B cell transcription factor Pax5 locus (Left). HiChIP - Hi-C delta heat map at the Pax5 locus (Right). **e,** Virtual 4C interaction profile at the XIST promoter in GM12878 for *in situ* Hi-C and HiChIP. **f,** Meta analysis of signal enrichment, scaled for interaction distance, at a union set of loops in Smc1a HiChIP and *in situ* Hi-C.

We then sought to characterize our high-confidence loop calls made with Juicer in GM Smc1a HiChIP. We investigated cohesin and CTCF binding at both ends of our loops and found that 81% of our loops were anchored by cohesin and 80% by CTCF, in close agreement with *in-situ* Hi-C data^5^**(Supplementary Fig. 2b).** In addition, our confident loop set exhibited a similar CTCF motif orientation distribution as compared to *in-situ* Hi-C^5^, with 80% of our loops exhibiting convergent CTCF motifs **(Supplementary Fig. 2c).** Furthermore, our contact signal was highly reproducible across biological replicates **(Supplementary Fig. 3).** We next examined the reproducibility of our low cell number HiChIP and found the 10, 5, and 1M experiments retained strong correlation to the 25M cell HiChIP **(Supplementary Fig. 4).** Finally, we assessed the characteristics of the non-overlapping *in situ* Hi-C and HiChIP loops (Fig. 2b). By examining CTCF and cohesin binding at the loop ends we found similar recruitment, suggesting that these classes are in fact biologically similar to one another **(Supplementary Fig. 5b).**

To better understand how HiChIP achieved similar loop calls from less sequencing, we investigated the nature of the contact map enrichment in Smc1a HiChIP. We first generated 5 kilobase resolution read-normalized interaction maps in both Hi-C and HiChIP, and then subtracted the Hi-C signal from the HiChIP signal to generate a differential heat map. We observed enrichment of chromatin loops in HiChIP and depletion elsewhere in the matrix (Fig. 2d). Our findings are consistent with the fact that the majority of *in situ* Hi-C identified loops are anchored with cohesin and CTCF^5^. Therefore, performing affinity-based pull-down of interactions associated with cohesin (or in principle CTCF) via HiChIP will decrease background and increased signal of long-range contacts. The increased enrichment for contacts was apparent at numerous other loci in both GM and mES HiChIP data, including in our low cell number samples **(Supplementary Fig. 6).**

To precisely visualize enrichment of HiChIP relative to *in situ* Hi-C we employed virtual 4C, where a specific genomic region is selected as an anchor “view point”, and all PETs connecting to that anchor are visualized as a line plot. Anchoring our analysis at a cohesin peak near the XIST RNA promoter, we first looked at the X-chromosome inactivation center (XIC) as this locus has been extensively characterized by multiple orthogonal conformation techniques and imaging^5,22,23^. HiChIP signal was strongly correlated with *in situ* Hi-C signal and exhibited a higher signal to background ratio (Fig. 2e).

For a global comparison of signal to background in HiChIP as compared to *in situ* Hi-C, we plotted the 4C signal profiles of the union set of loops from each experiment as heat maps sorted by the distance between the two loop ends and centered on the downstream loop anchor **(Supplementary Fig. 7).** We observed that HiChIP exhibited increased signal at the loop anchors as well as within the loops **(Supplementary Fig. 7).** We examined this quantitatively by performing meta-analysis of the signal enrichment, scaled for the loop interaction distance **(Methods).** HiChIP generated higher signal at chromatin loops relative to local background, as compared to *in situ* Hi-C (Fig. 2f).

To assess the general applicability of HiChIP, we performed the method targeting Oct4 in mES cells. This transcription factor has not been examined by ChIA-PET and can be compared with our Smc1a mES cell dataset. We found that Oct4 loops were largely observed in the Smc1a dataset, while Smc1a was also associated with loops exhibiting minimal Oct4 signal **(Supplementary Fig. 8).** Loops with high Oct4 contact signal relative to Smc1a were enriched for overlap with RNA polymerase II and depleted for overlap with CTCF, consistent with the notion that joint Oct4-Smc1a loops are associated with active enhancers and promoters^13,24^.

We also considered the potential for bias in HiChIP signal due to the enrichment of target-bound loci. The two contact calling tools that we implemented, Fit-Hi-C and Juicer, rely on coverage normalization and KR matrix balancing, respectively, to correct for potentially non-biological signal observed in regions that appear highly visible in a Hi-C-like experiment^5,19–21^. With and without normalization, we compared HiChIP signal at observed loops as well as random pairs of cohesin-bound loop ends to represent a background **(Supplementary Fig. 9).** We conclude that either normalization, while not developed to address HiChIP data specifically, appears effective in minimizing background signal while retaining robust signal enrichment at true loops. Further development of HiChIP-specific analysis tools may be useful for background removal.

Here we present HiChIP, a rapid, efficient, and technically simplified way to assay protein-centric chromatin conformation. By enriching for cohesin, we showed that HiChIP provides an efficient alternative strategy to deeply sequenced *in situ* Hi-C. Finally, the ability to sustain high confidence contact maps at low cell number will facilitate the investigation of chromatin conformation in systems previously un-measurable by conventional strategies.

## Accession Codes

NCBI Gene Expression Omnibus: raw and processed data available at accession number GSE80820.

## Acknowledgments

We thank our lab members for discussion. We thank N. Suliman for her critical reading of the manuscript. Supported by the Stanford Genome Training Program (NIH/NHGRI) (M.R.M.), Human Frontier Science Program, Rita Allen Foundation, Bio-X Stanford Interdisciplinary Graduate Fellowship (A.J.R), National Institutes of Health (NIH) 1F30CA189514-01 Stanford Medical Scientist Program (R.A.F.), NIH P50-HG007735 (to. H.Y.C., W.J.G.), and NIH S10OD018220 to Stanford Functional Genomics Facility.

## Author Contributions

M.R.M and R.A.F developed the method. M.R.M performed experiments. A.J.R and C.D analyzed the data. M.R.M, A.J.R, R.A.F, C.D, P.A.K, W.J.G, and H.Y.C interpreted the results and wrote the manuscript.

## Methods

### Cell culture and fixation

GM12878 cells (Coriell) were grown in RPMI 1640 (Gibco) with 15% FBS to a concentration of 500,000 to 1 million cells per mL. Mouse ES cells (V6.5 from Novus Biologicals: NBP1-41162) were grown in Knockout DMEM (Gibco) with 15% FBS and LIF (Millipore) to about 80% confluence. Detached adherent or suspension cells were then pelleted and resuspended in freshly made 1% formaldehyde (Thermo Fisher) at a volume of 1 mL of formaldehyde for every one million cells. Cells were incubated at room temperature for 10 minutes with rotation. Glycine was then added to a final concentration of 125 mM to quench the formaldehyde. Cells were incubated at room temperature for 5 minutes with rotation. Cells were pelleted and washed in PBS, then pelleted again and stored at −80° C or immediately taken into the HiChIP protocol.

### HiChIP

#### In situ Contact Generation

*In situ* contact libraries were generated according to the *in situ* Hi-C published protocol^5^ through the proximity ligation with modifications. In brief, up to 15 million crosslinked cells were resuspended in 500 μL of ice-cold Hi-C lysis buffer (10 mM Tris-HCl pH 7.5, 10 mM NaCl, 0.2%NP-40, 1X Roche protease inhibitors −11697498001) and rotated at 4° C for 30 minutes. For cell amounts greater than 15 million, the cell pellet was split in half for contact generation and then recombined for the sonication. Nuclei were pelleted at 4° C for 5 minutes at 2500 rcf and the supernatant was discarded. Pelleted nuclei were washed once with 500 μL of ice-cold Hi-C lysis buffer. Supernatant was removed again and pellet was resuspended in 100 μL of 0.5% SDS and incubated at 62° C for 10 minutes with no shaking or rotation. 285 μL of water and 50 μL of 10%Triton X-100 were added and samples were rotated at 37° C for 15 minutes to quench the SDS. 50 μL of NEB Buffer 2 and 15 μL of 25 U / μL MboI restriction enzyme (NEB, R0147) were then added and sample was rotated at 37° C for 2 hours. For lower starting material less restriction enzyme was used: 15 μL was used for 10-15 million cells, 8 μL for 5 million cells, and 4 μL for 1 million cells. MboI was then heat inactivated at 62° C for 20 minutes with no shaking or rotation. To fill in the restriction fragment overhangs and mark the DNA ends with biotin, 52 μL of incorporation master mix was then added: 37.5 μL of 0.4 mM biotin-dATP (Thermo Fisher 19524016), 4.5 μL of a dCTP, dGTP, and dTTP mix at 10 mM each, and 10 μL of 5 U / μL DNA Polymerase I, Large (Klenow) Fragment (NEB, M0210). The reactions were then rotated at 37° C for 1 hour. 948 μL of ligation master mix was then added: 150 μL of 10X NEB T4 DNA ligase buffer with 10 mM ATP (NEB, B0202), 125 μL of 10% Triton X-100, 3 μL of 50 mg/mL BSA (Thermo Fisher AM2616), 10 μL of 400 U / μL T4 DNA Ligase (NEB, M0202), and 660 μL of water. The reactions were then rotated at room temperature for 4 hours. After proximity ligation, the nuclei with in-situ generated contacts were pelleted at 2500 rcf for 5 minutes at room temperature and the supernatant was removed.

#### Sonication and Chromatin Immunoprecipitation

The nuclear pellet was brought up to 880 μL in Nuclear Lysis Buffer (50 mM Tris-HCl pH 7.5, 10 mM EDTA, 1% SDS, 1X Roche protease inhibitors −11697498001) and transferred to a Covaris millitube. Samples were sheared using a Covaris E220 using the following parameters: Fill Level = 10, Duty Cycle = 5, PIP = 140, Cycles/Burst = 200, Time = 4 minutes and then clarified by centrifugation for 15 minutes at 16100 rcf at 4° C. We kept the sonication constant at 4 minutes for different amounts of cell starting material, although sonication time may need to be adjusted for different cell types. The ideal sonication time will be as short as possible to allow for efficient ChIP signal over background. Too long of a sonication can lead to separation of the protein factor from the biotin contact, which will increase the likelihood of a DNA fragment not making it through both enrichments and a loss in sample complexity. Clarified samples were transferred to eppendorf tubes and 2X volume of ChIP Dilution Buffer (0.01% SDS, 1.1% Triton X-100, 1.2 mM EDTA, 16.7 mM Tris-HCl pH 7.5,167 mM NaCl) was added (samples were split between two eppendorf tubes to allow for the volume to fit at approximately 1.2 mL in each tube with a total of 2.4 mL for the entire ChIP). For our Smc1a (Bethyl A300-055A) and Oct4 (Santa Cruz 8628) antibodies we dilute 1:2 in ChIP Dilution Buffer to achieve a SDS concentration of 0.33%. However, for other antibodies a lower SDS amount may be necessary, and lysate can be diluted up to 1:9 to achieve a SDS concentration of 0.1%. For HiChIP of Smc1a in 10 million cells we first precleared with 60 μL of Protein A beads (Thermo Fisher). Beads were washed and resuspended in ChIP Dilution Buffer to a volume of 50 μL per tube (100 μL per HiChIP), then added to samples and rotated at 4° C for 1 hour. Samples were placed on magnet and supernatants were transferred into fresh tubes. For HiChIP of Smc1a in 10 million cells we added 7.5 μg of antibody and incubated overnight at 4° C. We then captured with 60 μL of Protein A beads that were washed and resuspended in ChIP Dilution Buffer to a volume of 50 μL per tube (100 μL per HiChIP). Beads were added to sample, and rotated at 4° C for 2 hours. Amounts of beads (for preclearing and capture) and antibody were adjusted linearly for different amounts of cell starting material. For 25 million cells we used 18.75 μg of antibody and 150 μL of beads, for 5 million cells we used 3.75 μg of antibody and 30 μL of beads, and for one million cells we used 0.75 μg of antibody and 6 μL of beads. For Oct4 we used the same amounts and conditions as Smc1a, except we used Protein G beads rather than Protein A for both preclearing and capturing. Furthermore, while these conditions given are what were used for Smc1a and Oct4, other factors may require different amounts or conditions. After bead capture, beads were washed three times each with Low Salt Wash Buffer (0.1% SDS, 1% Triton X-100, 2 mM EDTA, 20 mM Tris-HCl pH 7.5, 150 mM NaCl), High Salt Wash Buffer (0.1% SDS, 1% Triton X-100, 2 mM EDTA, 20 mM Tris-HCl pH 7.5, 500 mM NaCl), and LiCl Wash Buffer (10 mM Tris-HCl pH 7.5, 250 mM LiCl, 1% NP-40, 1% sodium deoxycholate, 1 mM EDTA, make fresh). Washing was performed at room temperature on a magnet by adding 500 μL of a wash buffer, swishing the beads back and forth twice by moving the sample relative to the magnet, and then removing the supernatant.

#### DNA Elution and Reverse Crosslinking

Sample beads were resuspended in 150 μL of DNA Elution Buffer (50 mM sodium bicarbonate pH 8.0, 1% SDS, make fresh) and incubated at room temperature for 10 minutes with rotation, followed by 3 minutes at 37° C shaking. Beads were then placed on a magnet and supernatant was transferred to a fresh tube. Another 150 μL of DNA Elution Buffer was added to beads and incubations were repeated. Supernatant was removed again, and 15 μL of Proteinase K (Thermo Fisher) added to the 300 μL reaction. Samples were incubated at 55° C for 45 minutes with shaking then temperature was increased to 67° C for 1.5 hours with shaking. Samples were purified with DNA Clean and Concentrator columns (Zymo Research) and eluted in 10 μL of water (elutions were done with the same 10 μL of water twice to achieve a higher recovery of DNA).

#### Biotin Capture and Preparation for Illumina Sequencing

Post-ChIP DNA was quantified by Qubit (Thermo Fisher) to estimate the amount of Tn5 (Illumina) needed to generate libraries at the correct size distribution (this is assuming contact libraries were generated properly, samples were not oversonicated, and material will robustly capture on streptavidin beads). For libraries with greater than 150 ng of post-ChIP DNA, material was set aside and a maximum of 150 ng was taken into the biotin capture step. For Smc1a HiChIP with 10 million cells an expected yield of post-ChIP DNA can be anywhere from 15 ng to 50 ng depending on the cell type. 5 μL of Streptavidin C-1 beads (Thermo Fisher) were washed with Tween Wash Buffer (5 mM Tris-HCl pH 7.5, 0.5 mM EDTA, 1 M NaCl, 0.05% Tween-20) then resuspended in 10 μL of 2X Biotin Binding Buffer (10 mM Tris-HCl pH 7.5, 1 mM EDTA, 2M NaCl). Beads were added to the samples and incubated at room temperature for 15 minutes with shaking. After capture, beads were placed on a magnet and supernatant was discarded. Samples were washed twice by adding 500 μL of Tween Wash Buffer and incubated at 55° C for 2 minutes shaking. Samples were then washed in 100 μL of 1X TD Buffer (2X TD Buffer is 20 mM Tris-HCl pH 7.5, 10 mM magnesium chloride, 20% dimethylformamide). After washes, beads were resuspended in 25 μL of 2X TD Buffer, Tn5 (for 50 ng of post-ChIP DNA we used 2.5 μL of Tn5), and water to 50 μL. Tn5 amount was adjusted linearly for different amounts of post-ChIP DNA, with a maximum amount of 4 μL of Tn5. For example 25 ng of DNA was transposed using 1.25 μL of Tn5, while 125 ng of DNA was transposed with 4 μL of Tn5. Using the correct amount of Tn5 is critical to the HiChIP protocol to achieve an ideal size distribution. An overtransposed sample will have shorter fragments and will exhibit lower alignment rates (when the junction is close to a fragment end). An undertransposed sample will have fragments that are too large to cluster properly on an Illumina sequencer. A maximum amount of Tn5 is used in order to save on Tn5 costs, and considering that a library with this much material will be amplified in 5 cycles and have enough complexity to be sequenced deeply regardless of how fully transposed the library is to achieve an ideal size distribution. Samples were incubated at 55° C with interval shaking for 10 minutes. Beads were then placed on a magnet and supernatant was removed. 50 mM EDTA was added to samples and incubated with interval shaking at 50° C for 30 minutes. Samples were then placed on a magnet and supernatant was removed. Samples were washed two times each in 50 mM EDTA then Tween Wash Buffer at 55° C for 2 minutes. Beads were lastly washed in 10 mM Tris before PCR amplification.

#### PCR and Size Selection

Beads were resuspended in 25 μL of Phusion HF 2X (New England Biosciences), 1 μL of each Nextera Ad1_noMX and Nextera Ad2.X at 12.5 μM^25^(Supplementary Table 4), and 23 μL of water. The following PCR program was performed: 72° C for 5 minutes, 98° C for 1 minute, then cycle at 98° C for 15 seconds, 63° C for 30 seconds, and 72° C for 1 minute. Cycle number was estimated using one of two different methods: (1) Reactions were first run 5 cycles on a regular PCR and then removed from beads. 0.25X SYBR green was added and then run on a qPCR where samples were pulled out at the beginning of exponential amplification. (2) Reactions were run on a PCR and cycle number was estimated based on the amount of material from the post-ChIP Qubit (approximately 50 ng was ran in six cycles, while 25 ng was ran in seven, 12.5 ng was ran in eight, etc.). The 1m cell samples had 2 ng of post-ChIP DNA, were transposed with 0.1 μL of Tn5 (1 μL of a 1:10 dilution), and were amplified in 10 cycles.

For one million cell samples reactions were PCR amplified for 5 cycles then placed on a magnet and eluted into new tubes. Libraries were then pooled and primers were removed with an Ampure XP cleanup (followed standard protocol as per manufacture, Beckman Coulter). Samples were further amplified five cycles (98° C for 15 seconds, 55° C for 30 seconds, and 72° C for 1 minute) with short primers. Pooling samples was done to increase likelihood of library amplification over primer artifact amplification. However, this is not an essential step and separate amplification for one million cell samples still generates high complexity libraries. The short primer sequences are:

Nextera_i7Short = CAAGCAGAAGACGGCATACGAGAT

Nextera_i5Short = AATGATACGGCGACCACCGAGATCTACAC

Size selection was performed using one of two different methods: (1) PAGE size selection of the final library. After PCR libraries were placed on a magnet and eluted into new tubes then purified with DNA Clean and Concentrator columns to a volume of 10 μL. Amplified DNA was run out on a 6% PAGE gel, stained with SYBR, and cut to a size range of 300-700. Note that if the bulk of the material is smaller those sizes can be included but the paired-end libraries will have a lower alignment rate. In the future Tn5 amount was adjusted accordingly. (2) Two-sided size selection with the Ampure XP beads. After PCR libraries were placed on a magnet and eluted into new tubes. 25 μL of Ampure XP beads were added and the supernatant was kept to capture fragments less than 700 bp. Supernatant was transferred to a new tube and 15 μL of fresh beads were added to capture fragments greater than 300 bp. After size selection, libraries were quantified with qPCR against Illumina primers and/or Bioanalyzer. Libraries were sequenced paired-end with read lengths of 75.

### HiChIP Data Processing

HiChIP paired-end reads were aligned to hg19 or mm9 genomes using the HiC-Pro pipeline^16^. Default settings were used to remove duplicate reads, assign reads to MboI restriction fragments, filter for valid interactions, and generate binned interaction matrices.

### HiChIP Contact Calling with Fit-HiC, Mango, and Juicer

HiCCUPS will not run on an interaction matrix that is too sparse (less than 300m filtered reads), and so for our low cell number samples which were not sequenced as deeply, we used the Fit-HiC contact caller. We also processed our 25m cell libraries with Fit-HiC for the comparisons in Figure 1d and Supplementary Figure 2a.

Bin pairs of the interaction matrix with statistically significant contact signal were identified using Fit-HiC^19^. Genome wide intra-chromosomal bin pairs were filtered for an interaction distance between 20kb and 2Mb, and default Fit-HiC settings were used to calculate false discovery rate (FDR) values for each bin-pair in a given HiChIP experiment. For the comparison of mESC experiments in Figure 1D, Smc1a HiChIP bin-pairs or Smc1a ChIA-PET high-confidence interactions^13^ were filtered for overlap with Smc1a ChIP-seq peaks. The appropriate FDR was selected for each HiChIP experiment to result in approximately 10,000 contacts per experiment.

GM12878 Smc1a HiChIP filtered PETs from the HiC-Pro pipeline were processed through stages 4 and 5 of the Mango^17^ pipeline to call significant interactions. Stage 4 applies MACS2^18^ to call peaks using PETs. Stage 5 modeled the background interactions by taking interaction distance and depth into consideration and used a binomial distribution to call significant interactions. With our Mango parameters of PETs >= 4 and a FDR cutoff of 10^-4^ we obtained 61395 significant interactions.

The Juicer pipeline's HiCCUPS tool was used to identify loops^5,20,21^. Filtered read-pairs from the HiC-Pro pipeline were converted into .hic format files and input into HiCCUPS.

The same parameters used for GM12878 Hi-C^5,20,21^ were used on the HiChIP datasets as follows: hiccups -m 500 -r 5000,10000 -f 0.1,0.1 -p 4,2 -i 7,5 -d 20000,20000 HiCCUPS_output.txt

### HiChIP ChIP Peak Calling

Dangling-end and self-ligation reads from the GM12878 Smc1a HiChIP HiC-Pro output were combined and processed to be compatible as a MACS2 BED file input. MACS2 was then run on the BED file using the no model and extsize 147 parameters and a FDR cutoff of 1%. Smc3 ChlP-seq peak sets for GM12878 were downloaded from the UCSC ENCODE repository. Bedtools intersect was run on Smc3 ChIP peaks and Smc1a HiChIP peaks to determine overlap. The bedtools intersect -v option was used to find peaks that did not overlap, and MACS2 -log10 q values were plotted for HiChIP overlapping and non-overlapping peaks.

### Loop Overlap Statistics for Venn Diagrams

The overlap and unique sets of loops were identified by comparing loop ends for all pairs of loops. For a given pair of loops, if both pairs of loop ends were separated by at most one bin, the loops were called as overlapping. The set of loops remaining as non-overlapping were considered unique to one loop set.

### CTCF Motif Orientation Analysis

The genome-wide set of motifs was obtained from the ENCODE motif repository^26^. Sets of contacts were first filtered for contacts in which each contact end overlapped only a single CTCF motif match. The proportion of contacts with convergent (+ strand motif on upstream end /- strand motif on downstream end) divergent (-/+), or in the same orientation (+/+ or -/-) were counted and divided by the total number of contacts overlapping a single motif on each end.

### Encode TF ChIP-seq Peak Overlap Enrichment in Juicer Loops

Comprehensive analysis of transcription factor ChIP-seq binding breadth and enrichment loop ends in GM12878 **(Supplementary Fig. 2D)** was performed as previously described^5^. The Smc3 and CTCF ChIP-seq peak sets corresponding to GM12878 were downloaded from the UCSC ENCODE repository. For each peak set, the proportion of peaks overlapping loop ends and randomly shuffled loop ends were determined. The proportion of peaks overlapping loop ends was divided by the proportion of peaks overlapping shuffled loop ends for each peak set. The resulting enrichment values were divided by the average enrichment for all peak sets and reported as the relative enrichment.

### Interaction Matrices and Virtual 4C Visualization

Hi-C and HiChIP heat maps presenting all filtered reads were generated with Juicebox^5,20,21^. For HiChIP matrices, .hic format files were generated as described above.

Depth-normalized heat maps of Hi-C or HiChIP interaction matrices were visualized as heat map using the heatmap.2 function from the gplots package in R. Hi-C interaction raw matrices were taken from the GEO repository GSE63525. HiChIP interaction matrices were generated with HiC-Pro as described above. Virtual 4C plots were generated from Hi-C or HiChIP interaction matrices by filtering the matrix for all bin-pairs in which one bin matched a single anchor bin. For virtual 4C or heat maps, depth-normalization was achieved by scaling counts by the total number of filtered reads in each experiment. The delta heat map was generated by subtracting the depth-normalized Hi-C matrix from the depth-normalized HiChIP matrix.

### Reproducibility Scatter Plots and Correlations

Experimental reproducibility scatter plots and comparisons between Hi-C and HiChIP were generated by counting reads supporting a set of loop calls. For HiChIP replicate comparisons in GM12878 or mESC, the corresponding loops identified from the 25m cell merged dataset was used. For the comparison between HiChIP and Hi-C, a union loop set between Hi-C and HiChIP was generated by merging loops identified in the HiChIP 25m cell merged dataset and the Hi-C primary + replicate^5^, followed by collapsing exact overlap loops.

The pearson correlation between replicates or experiments was calculated from raw reads using the cor() function in R. For generating scatter plots, reads were depth-normalized to 10m filtered fragments and then quantile normalized across the pair of replicates or experiments under consideration.

### Loop 4C Heat Maps and Loop Distance-Scaled Metaplots

Virtual 4C heat maps were generated by first making profiles from the upstream end of each loop in a set. These profiles were then sorted by end-to-end loop distance and aligned to the downstream anchor. Loops with a distance less than 1Mb only were presented to facilitate visualization. Distance-scaled metaplots were generated by dividing the locus containing a loop into three sections: upstream of the loop, intra-loop, and downstream of the loop. The intra-loop section was divided into 200 equally spaced bins. The upstream section was composed of 50 bins of the same size as the intra-loop bins, and the downstream section was composed of 100 bins of the same size as the intra-loop bins. The resulting 350 bin profiles were then combined into a matrix, and bin signal averages were calculated to form a metaplot.

### Comparison of PoI2 and CTCF enrichment at Oct4 and Smc1a-biased loops

Oct4 and Smc1a-biased loops were identified by comparing the depth normalized reads of Oct4 and Smc1a signal in the union set of loops identified from HiChIP of each factor. The top 500 biased loops out the union set of 11,035 were analyzed for overlap of Pol2 and CTCF, and the Oct4/Smc1a ratio of the proportion of Pol2 or CTCF overlapping loop anchors is presented.

### Comparison of normalization strategies

Metaplots of raw counts, VC-normalized, and KR-normalized GM12878 Smc1a HiChIP signal at loops ends were generated using contact matrices exported from Juicer^5,20,21^. Loops were filtered for the subset in which a cohesin (Smc3) ChIP-seq peak was identified at each loop end. Random pairs of anchors containing cohesin ChIP-seq peaks were selected to match the distance distribution of the observed loops as a reference for expected background signal.

